# Stuck in the weeds: Invasive grasses reduce tiger snake movement

**DOI:** 10.1101/2023.03.06.531246

**Authors:** Jari Cornelis, Christine E Cooper, Damian C Lettoof, Martin Mayer, Benjamin M Marshall

## Abstract

Wetlands are particularly vulnerable to degradation in urban environments, partially due to the introduction of non-native plants. Invasive weeds in wetlands can replace native plants leading to alterations in habitat composition and vegetation, in turn, animal movements and ultimately population dynamics might be affected. Here we investigate how home range size and movements of western tiger snakes (*Notechis scutatus occidentalis*) differ in wetlands dominated by invasive kikuyu grass (*Cenchrus clandestinus*) compared to wetlands dominated by native vegetation to understand if and how the movement ecology of this top-order predator is altered by vegetation homogenization. To do so, we used Autocorrelated Kernel Density Estimators (AKDE) to estimate home range size, dynamic Brownian Bridge Movement Models to document movement trajectory confidence areas, and compared movement distances using a Bayesian regression model. Home range sizes by tiger snakes were 14.59 ± 9.35 ha smaller in areas dominated by invasive versus native vegetation. Moreover, within-day movement distances tended to be smaller in areas dominated by invasive versus native vegetation (mean ± SD: 9 ± 3 m versus 18 ± 6 m), but there was considerable overlap between the 95% credible intervals between these two groups. Smaller home ranges by tiger snakes in areas dominated by invasive kikuyu grass were likely driven by thermoregulation, with snakes moving vertically between basking locations on top of kikuyu and shelter sites at the base, rather than travelling horizontally along the ground to open basking areas in sites dominated by native vegetation. Additionally, fragmentation of sites dominated by invasive vegetation might have contributed to the comparatively smaller home ranges of snakes there. These findings add to our understanding how changes in habitat composition driven by invasive vegetation can affect animal space use and emphasise the need for further studies to understand how these changes affect population dynamics.

## INTRODUCTION

Wetlands are sensitive ecosystems and are particularly vulnerable to degradation in urban environments (Faulkner 2004). Invasive weed species in wetlands (Grella et al. 2018) can replace native plant species, causing compositional and structural changes to the vegetation (Braithwaite et al. 1989; Reed et al. 2005). Habitat homogenisation by invasive plants can alter the structural complexity of microhabitats (Lambdon et al. 2008; Cornelis et al. 2022) and the availability of resources for animals (Schirmel et al. 2016) including the thermal quality of the vegetation (Hacking et al. 2015) and opportunities for camouflage (Valentine et al. 2007). Moreover, structural components of the vegetation can make mobility (Newbold 2005) and foraging (Maerz et al. 2005) more difficult, which can impact animal movement and behaviour (Doherty et al. 2019, Stewart et al. 2021). Consequently, invasive weeds can affect animal populations and communities. For example, habitat homogenisation by invasive cheatgrass (*Bromus tectorum*) in North America led to changes in small mammal community composition by increasing harvest mouse (*Rithrodontomys* spp.) and decreasing pocket mouse (*Perognathus* spp.) occupancy (Ceradini & Chalfoun 2017).

Species that rely on specific microhabitats and thermal conditions might be impacted most by the cumulative impacts of invasive plants (Devictor et al. 2008; Clavel et al. 2011). However, not all fauna are negatively affected by the invasion of exotic plants (Douglas et al. 2006). Some reptiles, for example, are more reliant on the structure provided by vegetation rather than the composition or species diversity, including invasive species, of the plant community (Garden et al. 2007; Hodgkinson et al. 2007; Garden et al. 2010). Some vegetation monocultures can provide favourable conditions for these species where they can persist, and even thrive, despite the predominance of a single invasive species (Lettoof et al 2021b). How animals react to differences in vegetation composition can be revealed through examining animal movement thereby providing an avenue to connect the impacts of vegetation and land use management decisions (Fraser et al., 2018). Consequently, investigating animal movement in weed-infested landscapes can lead to better informed management outcomes (Doherty and Driscoll 2018).

Western tiger snakes (*Notechis scutatus occidentalis*) persist in a handful of urban wetlands in Perth, Western Australia, a region where ∼70% of wetlands have been lost or degraded (Davis and Froend 1999; Kelobonye et al. 2019). The predatory role of tiger snakes in these wetlands (Lettoof et al. 2020) and evidence of their bioaccumulation of environmental contaminants make these snakes potentially useful bioindicators of urban wetland health (Lettoof et al. 2021). Many of Perth’s urban wetlands have lost their original riparian vegetation and are instead dominated by invasive flora (Davis and Froend 1999; Simpson and Newsome 2016) including kikuyu grass (*Cenchrus clandestinus*). Kikuyu grows as a dense matrix of stems, which facilitates its colonisation and can result in native plant communities being transformed into a monoculture of kikuyu (Gonzalez 2009; Bradshaw et al. 2013). The inter-plant distance of invasive grasses is often lower compared to native grasses, reducing the amount of bare ground that, together with limited plant diversity, results in reduced environmental heterogeneity (Litt and Steidl 2011; Lindsay and Cunningham 2012, Abom et al. 2015). Consequently, increased vegetation density and reduced availability of bare ground are two key structural features of kikuyu grass that differentiates it from the native riparian grass/tussock vegetation that occurs naturally in these wetlands (Cornelis et al. 2022).

Here we assess whether movements of western tiger snakes differ in wetlands dominated by invasive kikuyu grass compared to wetlands characterised by native vegetation. Our comparison aims to reveal if and how the movement ecology of this predatory species is altered by human-driven vegetation homogenization. We evaluated the potential effect of invasive kikuyu grass on tiger snake home range size (autocorrelated kernel density estimators; AKDE), movement trajectory confidence areas (dynamic Brownian bridge movement models; dBBMM), and compared step lengths per hour (using a Bayesian regression model) to gain insight into how tiger snakes may modify their movement in areas heavily affected by invasive vegetation.

## MATERIALS AND METHODS

### Study sites

We examined western tiger snake spatial ecology at four wetlands within 50km of the Perth CBD, Western Australia (Fig.1). Herdsman Lake (HS; 31.92°S, 115.80°E) and Kogolup Lake (KL; 32.12°S, 115.83°E) is dominated by invasive kikuyu grass. Loch McNess in Yanchep National Park (Y; 31.54°S, 115.68°E) and Black Swan Lake (BS; 32.47°S, 115.77°E) are dominated by native vegetation, predominantly *Schoenoplectus spp* in open habitats; *Ghania decomposita* and *Lepidosperma longitudinale* in sedgelands close to water; and *Banksia*, *Melaleuca* and *Eucalyptus spp* in woodlands (Cornelis et al. 2022; Fig. 1). For each site we measured the area (ha) of three land cover variables within 200m of the water’s edge (the greatest distance a snake travelled from the water) or less if there was a major barrier (e.g., building or roads) using QGIS v. 3.10.14 and ESRI satellite imaging after Lettoof et al. (2022). These variables were: total area, snake habitat (vegetation that contains mid-to-understory layers that tiger snakes could shelter in), and the percent of the snake habitat that was composed of kikuyu grass.

**Fig. 1.**
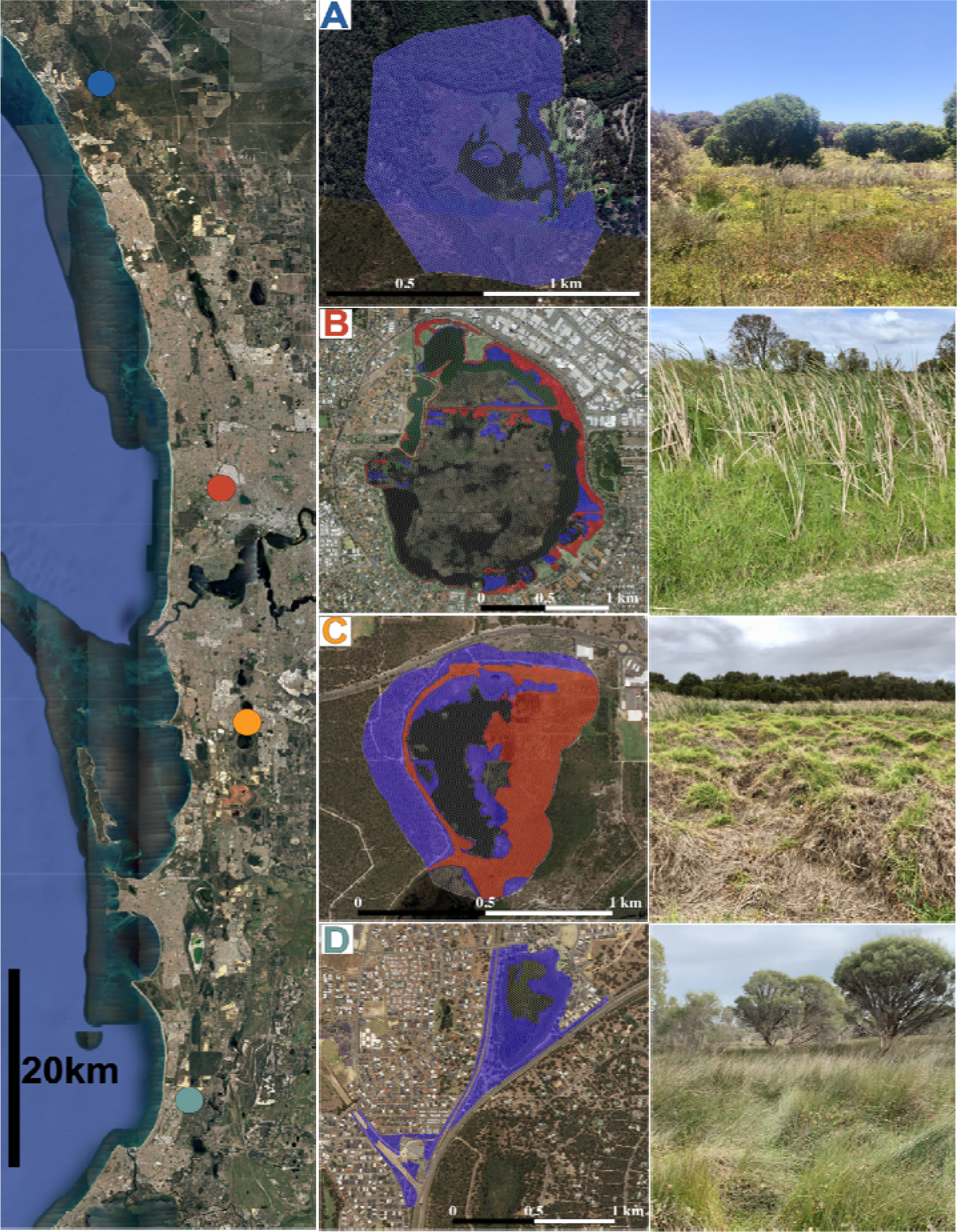
*Left panel*: The Perth metropolitan area, Western Australia with our four wetland *study* sites. Warmer colours (Herdsman Lake and Kogolup Lake) are sites dominated by invasive kikuyu grass (*Cenchrus clandestinus*) and cooler colours (Black Swan Lake and Yanchep National Park) are the sites with native vegetation. *Center panel*: Area of potential western tiger snake (*Notechis scutatus occidentalis*) habitat dominated by native (blue hashed) or introduced kikuyu grass (*Cenchrus clandestinus*; red hashed) at A) Yanchep National Park, B) Herdsman Lake, C) Kogolup Lake and D) Black Swan Lake. *Right Panel*: Representative habitat where tiger snakes spent the majority of their time during the tracking period corresponding to the site on the left of each image. Map Data: Imagery ©2022 Google, Imagery ©2022 CNES / Airbus, Maxar Technologies.

Of the total 322.5 ha area of the HL reserve an area of 49.1ha (15%) was potentially suitable for tiger snakes, and the remainder was mowed lawn or open water. Kikuyu grass dominated 61% of this potential tiger snake habitat, with the remainder native vegetation dominated by bulrush (*Typha sp*; Fig. 1). Kogolup Lake reserve total area was 68 ha area, with 54.2 ha (80%) potentially suitable for tiger snakes. Snakes were only caught around the northern half of Kogolup Lake, so we only measured this area of the wetland. Of this potential tiger snake habitat, 49% was dominated by kikuyu grass. The remaining native vegetation was dominated by *Eucalyptus*, *Melaleuca* and/or *Banksia spp* (Fig. 1).

There was no introduced kikuyu grass in potential tiger snake habitat at BS or Y lakes, with 40.5ha (69%) of the total 59 ha reserve native vegetation potentially suitable for tiger snakes at BS and 50.1 ha (80%) available of the total 62.8 ha area at Y (Fig. 1).

### Field methods: Capture, morphometrics and radio tracking

Adult male tiger snakes were hand-captured in September when they emerged from their overwinter- dormancy (Shine 1977; 1979). Only sexually mature male snakes (>650mm; Shine 1978) were studied as they are physically larger (to facilitate instrumentation) and are likely to be more active while searching for mates than females and juveniles (Shine 1979; Bonnet et al., 1999; Carfagno and Weatherhead, 2008). Their capture location was recorded with a Garmin (model 60) GPS. Snakes were weighed using a 500g Pesola spring balance, snout-vent length (SVL) and tail length were measured by stretching the snake along a ruler, and ventral scale clips were made for individual identification. The mean snout-vent-length of the 14 snakes was 84.6 ±1.51cm (range 72.8–96.0cm) and mass 294.46 ±15.21 g (207.5–427.5; Table 1). We ran ANOVAs on these measurements and determined that vegetation (native or kikuyu) had no effect on snake SVL or mass (p > 0.173). Sex was determined by inserting a lubricated probe into the cloacal bursae to measure the depth of the hemipenal pocket (McDiarmid et al. 2012). Fourteen snakes (Table 1) were transported to Curtin University campus where they were housed individually in plastic tubs (70x50x40cm) for up to a week prior to surgery, and then 2-5 days post-surgery to facilitate welfare monitoring. The snakes were not fed during the < two weeks they were in captivity, but fresh water was provided *ad lib*.

**Table 1.** Measurement and tracking data for individual tiger snakes (Notechis scutatus occidentalis) studied at four wetlands (HL, Herdsman Lake; KL, Kogolup Lake; BS, Black Swan Lake; Y, Yanchep National Park) in the Perth metropolitan area, Western Australia. A standard error accompanies mean values. Naive mean move distance is calculated using all step lengths, whereas daily mean move distance is calculated using step lengths where the tracking time lag was less than 24 hours.

| Snake ID | Snout-vent length (mm) | Mass (g) | Duration tracked (days) | Number of tracks | Median tracking time lag (hr) | Mean tracking time lag (hrs) | Number of moves | Naive mean move distance (m) | Daily mean move distance (m) |
| --- | --- | --- | --- | --- | --- | --- | --- | --- | --- |
| HL162 | 805 | 260 | 51 | 53 | 2.99 | 23.65 ± 4.98 | 52 | 18.8 ± 2.2 | 16.9 ± 2.6 |
| HL166 | 855 | 332.5 | 52 | 53 | 2.99 | 24.02 ± 5.06 | 52 | 10.9 ± 2.3 | 8.0 ± 2.1 |
| HL168 | 804 | 255 | 32 | 36 | 3.02 | 22.17 ± 5.96 | 35 | 5.2 ± 1.2 | 3.9 ± 0.4 |
| HL88 | 804 | 247.5 | 52 | 47 | 3.06 | 27.13 ± 5.69 | 46 | 51.8 ± 28.8 | 12.4 ± 2.4 |
| KL01 | 852 | 290 | 48 | 49 | 2.98 | 23.97 ± 5.27 | 48 | 17.8 ± 3.8 | 11.4 ± 3.2 |
| KL02 | 812 | 270 | 48 | 50 | 3.02 | 23.73 ± 5.16 | 49 | 12.0 ± 2.1 | 11.0 ± 2.4 |
| KL06 | 844 | 260 | 32 | 35 | 3.01 | 22.79 ± 6.12 | 34 | 6.5 ± 1.8 | 5.1 ± 2.0 |
| BS01 | 872 | 277.5 | 48 | 52 | 2.93 | 22.75 ± 5.02 | 51 | 62.9 ± 11.8 | 31.6 ± 6.0 |
| BS02 | 895 | 310 | 48 | 52 | 2.98 | 22.77 ± 5.01 | 51 | 12.9 ± 2.6 | 7.0 ± 1.5 |
| BS08 | 906 | 357.5 | 48 | 49 | 2.94 | 24.01 ± 5.29 | 48 | 92.9 ± 25.8 | 46.5 ± 16.2 |
| BS09 | 960 | 427.5 | 55 | 30 | 3 | 45.51 ± 18.9 | 29 | 97.6 ± 25.2 | 37.9 ± 9.1 |
| Y58 | 880 | 355 | 32 | 36 | 2.97 | 22.16 ± 6.01 | 35 | 12.8 ± 3.4 | 9.7 ± 1.5 |
| Y63 | 728 | 207.5 | 48 | 50 | 2.97 | 23.7 ± 5.23 | 49 | 25.5 ± 5.2 | 17.3 ± 3.8 |
| Y65 | 829 | 272.5 | 51 | 53 | 3 | 23.61 ± 5 | 52 | 34.35 ± 5.59 | 18.1 ± 2.9 |

A wax-coated (Elvax) VHF transmitter (Holohil PD-2; total mass ∼5g, <3% of the snakes’ body mass), was surgically implanted into the intraperitoneal cavity of each of the 14 snakes under general anaesthesia. Anaesthesia was induced by an intramuscular injection of Alfaxalone (5 mg kg^-1^) and maintained with gaseous isoflurane (1.5-4%) in oxygen. Local anaesthetic (lignocaine, 1 mg kg^-1^ and bupivacaine, 1 mg kg^-1^) was administered subcutaneously at the surgical site. The transmitter was inserted into the peritoneal cavity with the transmitter’s whip antenna inserted into a pocket under the skin. Analgesia was provided in the form of a subcutaneous injection of Meloxicam (0.2 mg kg^-1^).

Two to five days after surgery snakes were released at their point of capture. They were then radio-tracked for a maximum of two months during the period 12^th^ October 2020 to 3^rd^ December 2020. Due to the distance between study sites snakes could only be tracked at one site per day. We visited each site sequentially every four days and at each site we tracked individual snakes four times per day to determine their location, with approximately three hours between each location recording. Some snakes were collected before the two month period as their transmitter battery began to expire. Snakes were recaptured by hand and returned to Curtin University where they were euthanised via an intracardiac injection of Lethabarb (pentobarbitone 162.5 mg kg^-1^) and then dissected to remove the transmitter.

### Analysis: Space use estimates and comparing areas and movement

We estimated tiger snake home ranges using autocorrelated kernel density estimators (AKDE; Fleming & Calabrese, 2017), with the ctmm package (v.0.6.1; Calabrese et al. 2016) for R v.4.2.0. There are several advantages of AKDE over other estimators of home range (e.g., minimum convex polygon, kernel density estimators). Autocorrelated kernel density estimators use a fitted movement model to better estimate the potential locations an animal would travel, thereby fitting more closely to Burt’s (1943) definition of home range than movement-naïve traditional methods (Fleming & Calabrese, 2017). Autocorrelated kernel density estimators are also more robust for data with temporal gaps and they address the autocorrelation inherent in tracking data (Fleming et al. 2018; Noonan et al. 2019). We used perturbative hybrid REML (pHREML) to fit and determine the best fitting movement model for each individual –pHREML is well-suited for datasets with small effective sample sizes (Silva et al. 2021). We selected the model with the lowest AICc for each individual, and used the weighted 95% contour area estimates from that lowest AICc model in all further analysis. We used weighted estimates because of the gaps in data collection (Silva et al. 2021). Range stability, and therefore suitability for home range area estimation, was assessed with variograms. Values are presented as mean ± SD unless stated otherwise.

Autocorrelated kernel density estimators provide an estimate of animal’s home range, but due to the questionable stability of the ranges observed we supplemented these estimates with dynamic Brownian bridge movement models (dBBMM; Kranstauber et al., 2012; Kranstauber et al., 2022). Dynamic Brownian bridge movement models estimate the uncertainty surrounding movement pathways taken between known locations. The area estimates (or confidence areas) that dBBMM generate are considered an occurrence distribution (i.e., interpolation within a sampling period) that contrast with the use distributions provided by AKDEs (i.e., extrapolation to a full home range for an animal; Alston et al., 2022). Here we use dBBMM as a comparison of the potential areas the snakes could have reached between recorded locations. As we documented snake locations at similar frequency and durations, the differences in dBBMM confidence areas should reflect differences in movement rather than uncertainty derived from sampling variation (Silva et al., 2020) providing an additional line of evidence for any movement differences detected.

Dynamic Brownian bridge movement models base the estimates of uncertainty on the movement capacity of the animal (termed motion variance) calculated on a rolling basis from the tracking data (Kranstauber et al., 2012). The rolling basis is determined by two values, a window size and a margin size; we selected a large window (29 data points) and a margin likely still capable of detecting movement mode changes within that window (9 data points). We selected a broad window to mitigate the burst tracking regime, smoothing out spikes in movement activity that could be artefacts of the sampling protocol. We retrieved the areas from 90, 95, and 99% confidence area contours, and used the 95% contour in all further analyses. We used the R package move (v.4.1.8; Kranstauber et al., 2022) to run dBBMMs.

We compared the area estimates (home range from 95% AKDE and 95% confidence areas from dBBMM) using Bayesian comparative tests using a student distribution, as these provide more intuitive estimates of uncertainty with small samples (Morey et al., 2019). We used brms v.2.17.0 (Bürkner 2017, 2018, 2021), bayesplot v.1.9.0 (Gabry et al. 2019; Gabry and Mahr 2022), tidybayes v.3.0.2 (Kay 2022), and performance v.0.9.1 (Lüdecke et al. 2021), to run, visualise and explore these Bayesian models. For the Markov Chain Monte Carlo (MCMC) chains, we used 24,000 iterations, with a burn-in of 6,000, across 4 chains, and a thinning factor of 12. We used the resulting distributions to compare individuals at the sites with vegetation (*vegetation*) dominated by invasive grass sites (KL and HL) with those inhabiting sites dominated by native vegetation (Y and BS). We also included study site (*locale*) as a group effect to account for lack of independence between area estimates. The final formula was *area_estimate ∼ 0 + vegetation + (1|locale), sigma ∼ vegetation*. To avoid divergent transitions we increased adaptive delta to 0.9 and maximum tree depth to 15.

To explore whether the step lengths of individual movements (modelled using a lognormal distribution) differed based on whether they occupied an area with invasive or native vegetation we ran a Bayesian regression model. We included a binary population effect for *vegetation* (invasive versus native), and we included a nested group intercept effect to account for individual snakes (*snakeID*) and the study site they occurred in (*locale*). As location data were collected using a burst regime, we excluded all step lengths calculated between tracking days (i.e., no steps with a time lag greater than 24 hours were included). We accounted for the non-independence between data points collected on the same day by adding a second group intercept effect based on date. The final model used to explore invasive grass presence on step lengths per hour was: *step_length_over_hour ∼ 0 + vegetation + (1|locale/snakeID) + (1|date).* We ran the model in R using the brms package (Bürkner, 2017), using 8,000 iterations, 4 chains, with 4,000 burn-in, and a thinning factor of 2. To ensure convergence and to minimise divergent transitions (3 could not be prevented), we modified the maximum tree depth to 15 and the adaptive delta to 0.999. We used uniform priors, limited between 0 and 1,000, when exploring the difference between area estimates. We did not supply priors for the step length Bayesian models, applying the brm package defaults (flat priors for β degrees of freedom = 3, mu = 0, sigma = 1 for standard deviations). We determined chain convergence *R* using values (0.9 < *R* < 1.1) and reviewed trace, acf and posterior predictive check plots to check for other model convergence issues.

For the above analyses we used R v.4.2.0 (R Core Team 2022) via RStudio v.2022.7.1.554 (RStudio Team 2022). We used the tidyverse v.1.3.1 and reshape2 v.1.4.4 packages for data manipulation (Wickham et al. 2019; Wickham 2007). We used ggplot2 v.3.3.6 for creating figures (Wickham 2016), with the expansions: ggridges v.0.5.3 (Wilke 2021), ggpubr v.0.4.0 (Kassambara 2020), ggrepel v.0.9.1 (Slowikowski 2021), and ggspatial v.1.1.6 (Dunnington 2022). We used GADMTools v.3.9.1 (Decorps 2021), sp v.1.5.0 (Pebesma and Bivand 2005; R. S. Bivand et al., 2013), and rgeos v.0.5.9 (Bivand and Rundel 2021) to manipulate spatial data and plot country outlines. We used the ctmm v.0.6.1 to estimate AKDEs (Fleming and Calabrese 2021), and the move v.4.1.8 to estimate dBBMMs (Kranstauber et al., 2022). We used brms v.2.17.0 (Bürkner 2017, 2018, 2021), bayesplot v.1.9.0 (Gabry et al. 2019; Gabry and Mahr 2022), tidybayes v.3.0.2 (Kay 2022), and performance v.0.9.1 (Lüdecke et al. 2021), to run, visualise, and explore Bayesian models. We generated R package citations with the aid of grateful v.0.1.11 (Rodríguez-Sánchez et al. 2022).

## RESULTS

### Capture and tracking summary

We tracked snakes for a mean duration of 46.1 ±2.12 days (32–55; Fig. S1), collecting an average of 46.1 ±2.16 data points (30–53) per individual, and recording a mean of 45.1 ±2.16 moves per snake (29–52; Table 1). Overall, the burst sampling regime resulted in a mean time lag between data points of 24.6 ±1.65 hours (0.77–543.9 hours; Fig. S2), and a median time lag of 3 hours. Mean naive step length (including step lengths recorded between the bursts of sampling) was 32.3 ±3.49m (0.136– 1349m), and mean within-day step length (only steps recorded with a time lag of less than 24 hours) was 16.9 ±1.59m (0.26–542m).

### Area use and movement

At the two sites with native vegetation (BS and Y), we mostly observed tiger snakes in areas with native grasses (*Schoenoplectus* spp), whereas snakes were mostly observed in areas dominated by kikuyu grass in KL and HL. There was no evidence that the tiger snakes at KL and HL moved from their kikuyu sites to the adjacent native Banksia woodland during the period they were tracked. We frequently observed tiger snakes basking on top of the dense structure formed by kikuyu grass at both HL and KL while snakes at BS and Y were most commonly observed basking in sunny patches on the ground rather than on top of the vegetation (Fig. 2).

**Fig. 2.**
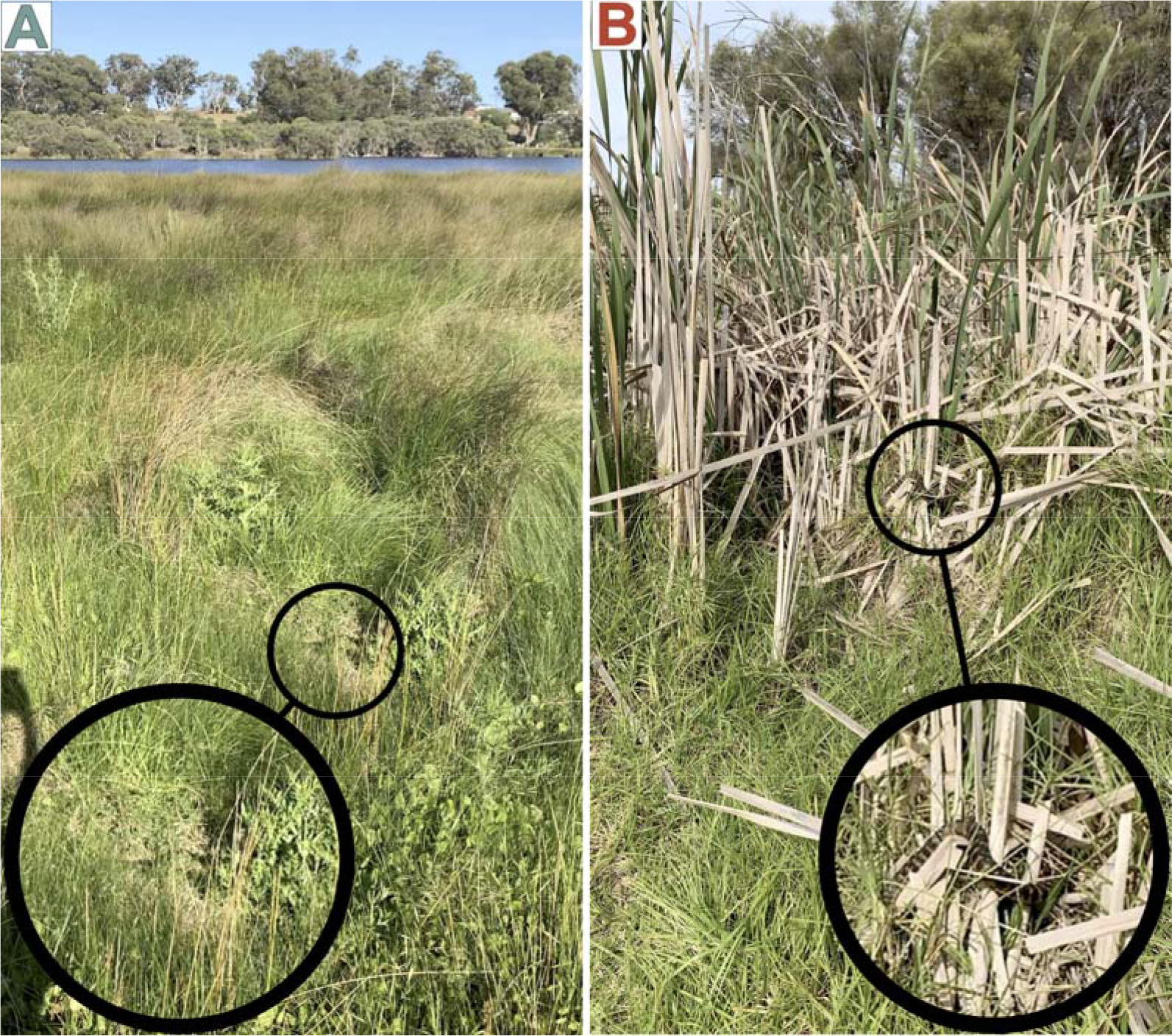
Two examples (including magnified insets in large circles) of how tiger snakes (*Notechis scutatus occidentalis*) were most commonly observed basking in (A) native vegetation at Black Swan Lake where the tiger snakes basked in sunny patches on the ground while in (B) invasive kikuyu grass at Herdsman Lake tiger snakes basked in elevated positions provided by the dense structure of kikuyu.

The 95% AKDE contours produced a mean estimated home range of 34.2 ±17.84 ha (range: 0.12–234.0 ha; Fig. 3). Invasive vegetation sites had a mean 95% point estimate of 35.85 ± 33.10 (range: 0.12 - 234.05 ha), whereas native vegetation sites had a mean 95% point estimate of 32.47 ±16.87 (range: 1.28 - 130.50 ha). However, the median range for invasive vegetation sites was 1.11 ha, compared to 21.5 ha at sites dominated by native vegetation. The Bayesian comparison indicated a 96.37% chance that AKDE home ranges (95% contour; and with the caveat of limited evidence of range residency) were smaller at the sites with invasive grass, with a mean difference of 14.6 ± 9.35 ha (Cr.I 95% -33.07–1.42 ha; Fig. 4A). The mean home range crossing time was 7.95 ±2.39 days for the snakes inhabiting sites with invasive vegetation compared to 5.18 ±1.96 days for those sites with native vegetation.

**Fig. 3.**
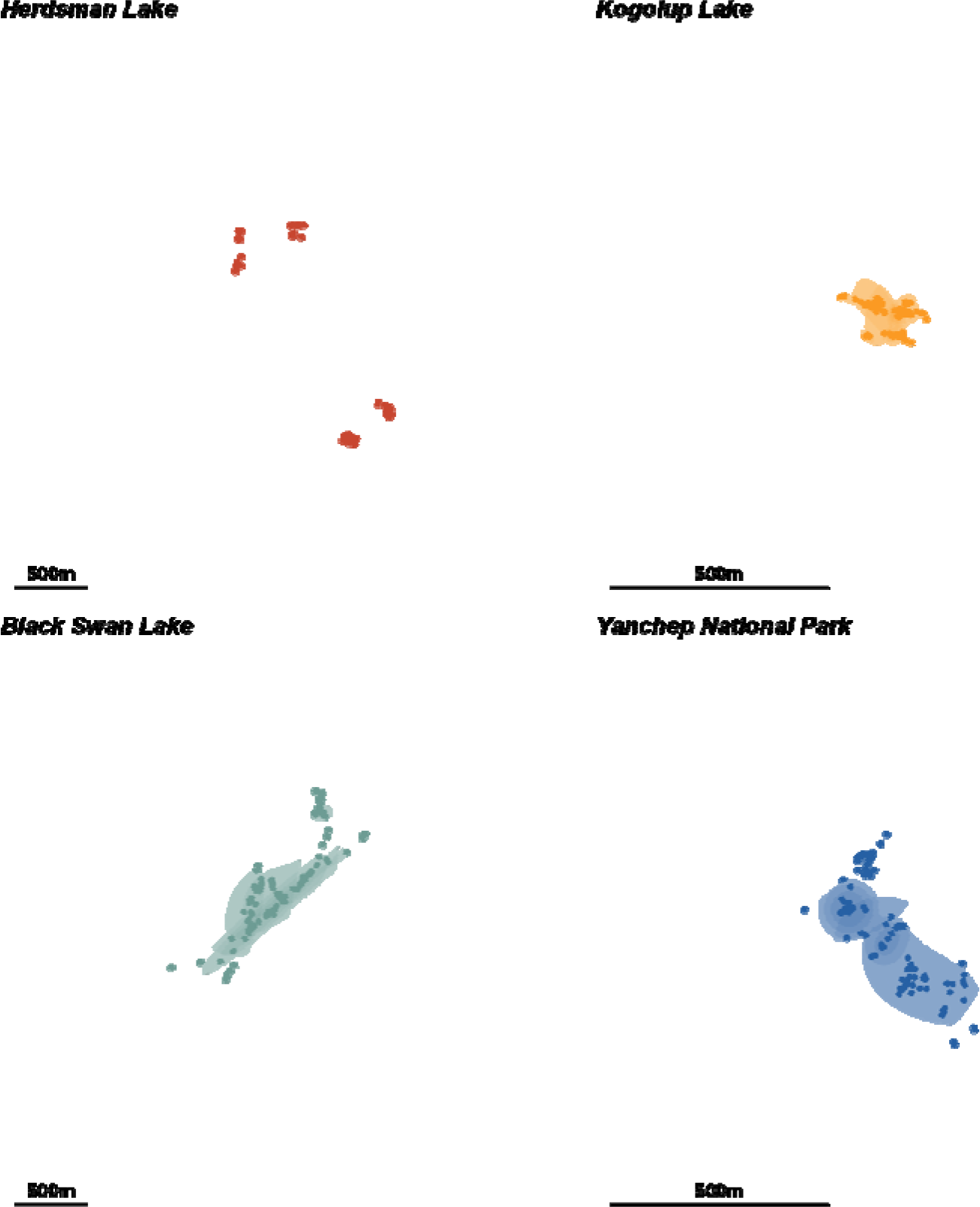
Autocorrelated Kernel Density Estimators (AKDE) area estimates for western tiger snakes (*Notechis scutatus occidentalis*) at four wetlands in the Perth metropolitan area, Western Australia. Warmer colours (Herdsman Lake and Kogolup Lake) are sites dominated by invasive kikuyu grass (*Cenchrus clandestinus*) and cooler colours (Black Swan Lake and Yanchep National Park) are the sites with native vegetation. Each snake’s home range area is represented by the 95% contour, alongside the 95% confidence surrounding that contour estimate in differing levels of transparency. Dots represent snake positions (raw data). The scale bar is 500 m.

**Fig. 4.**
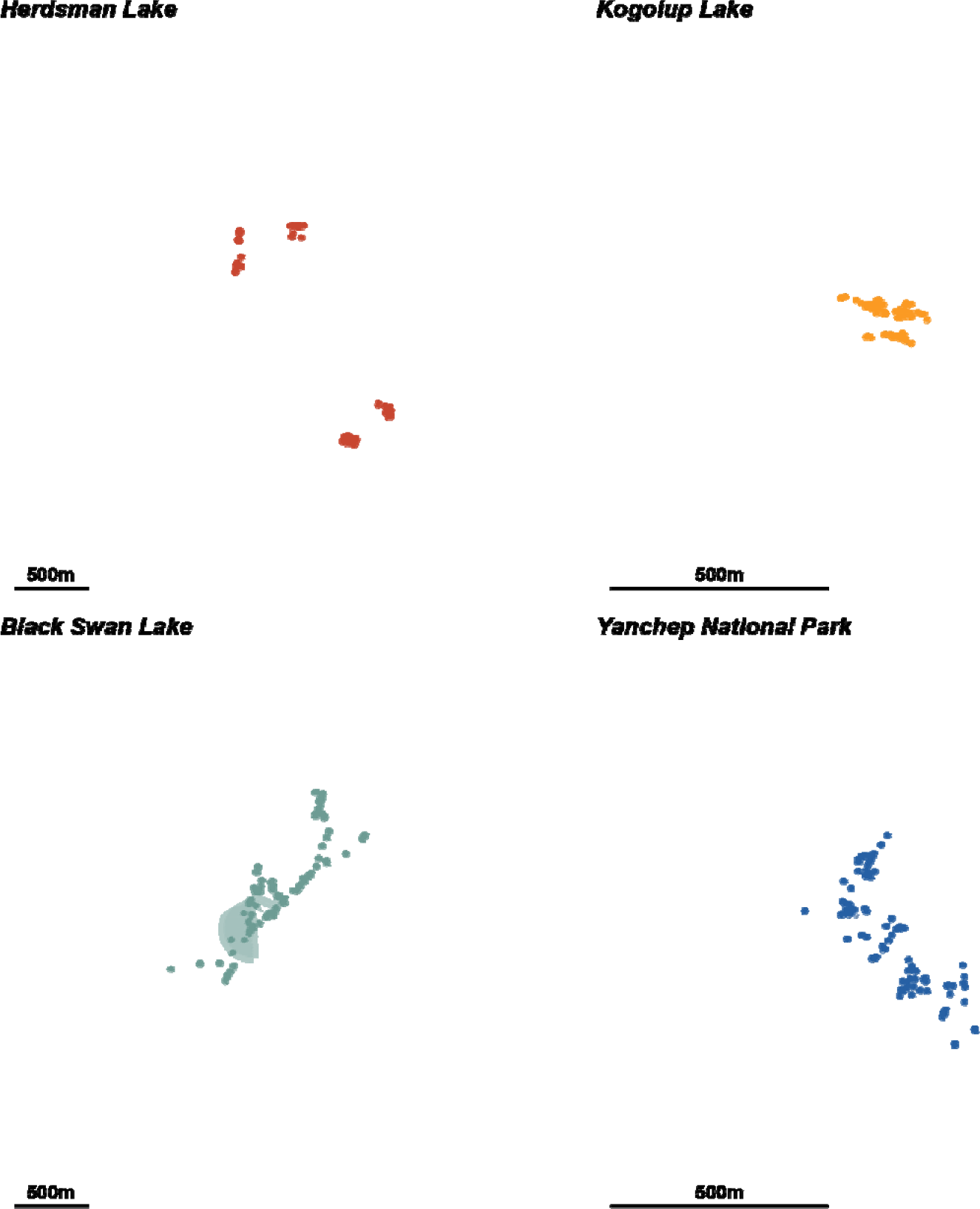
Dynamic Brownian Bridge Movement Models (dBBMM) confidence areas at each study locale for tiger snakes (*Notechis scutatus occidentalis*) radio-tracked at four wetlands in the Perth metropolitan area, Western Australia. Dots represent tiger snake positions (raw data). Scale bar is 500m, matching the scale and origin of the AKDE (Fig. 3) Warmer colours (Herdsman Lake and Kogolup Lake) are the locales with invasive grass; cooler colours (Black Swan Lake and Yanchep National Park) are the locales with native vegetation. Each snake’s confidence area is represented by the 90, 95 and 99% contours in differing levels of transparency.

Variograms for all snakes were severely impacted by the burst sampling tracking regime, with clear and repeated spikes in semi-variance uncertainty associated with the multi-day gaps (Fig. S2). Only snakes BS01 and BS02 had variograms that appeared to approach stability after peaks in semi- variance at time lags greater than 20-30 days. Stability of the other snakes’ ranges was difficult to ascertain because of the artefacts resulting from the burst sampling; therefore, range AKDE area estimates should be interpreted with caution (i.e., the assumption of range residency was not met). Lack of stability was compounded by low effective sample sizes for all individuals (10.9 ± 3.05, range: 1.34–37.80; Table S1), justifying the use of pHREML fitting and weighted area estimates (Silva et al., 2021). The lowest (0.08 ha) and highest (713.3 ha) 95% confidence intervals associated with the home range estimates illustrate the extent of uncertainty (Table S1) and the individual variation between individual snakes.

As the stability of the snakes’ range residency was uncertain (Fig. S3), we examined whether differences in potential space use are apparent using a different estimation method. Dynamic brownian bridge movement models (dBBMM; Fig. 4) provide an alternative estimate of uncertainty regarding possible areas reached by the snakes between data points. The broad window and margin sizes we selected successfully smoothed motion variance throughout the tracking period, avoiding artefact spikes in motion variance (movement capacity) resulting from the burst sampling (Fig. S4). The 95% confidence areas resulting from the dBBMMs revealed a clear difference in snake movements (i.e., areas potentially reached between recorded locations) between sites with native vegetation (11.68 ±4.80 ha, median: 4.97, range: 0.21–30.62 ha) and invasive vegetation dominated sites (0.57 ±0.43 ha, median: 0.08, range: 0.03–3.11 ha); snake movements at BS and Y areas (native vegetation) appeared much larger, implying greater uncertainty likely as a result of increased movement capacity detected by the models (Fig. 3; Table S1). Our Bayesian comparison reflected this difference, suggesting a 98.58% chance that space use was smaller in areas with invasive vegetation; on average 10.04 ±5.65 ha smaller (Cr.I 95% -20.52–0.14 ha; Fig. 4A).

The Bayesian Regression Model successfully converged with all *R* values ∼1, and trace and acf plots appeared adequate (see DOI: https://doi.org/10.5281/zenodo.7700983). However, a very low conditional R^2^ of 0.125 (Cr.I. 95% 0.065–0.213; marginal R^2^: 0.02, Cr.I. 95%1.127e-09–0.194), suggests factors other than those included in the model (binary vegetation categories or the group effects of animal, site and day) are impacting step length per hour. The mean (± SD) within-day step length per hour (i.e., those with a time lag of less than 24 hours) was 3.07 ±1.07 m/hour (Cr.I. 95% 0.81–10.31 m/hour) at sites with invasive vegetation compared to 6.19 ±2.04 m/hour (Cr.I. 95% 1.79– 20.36 m/hour; Fig. 5B) in areas with native vegetation. On average, the step lengths per hour recorded at with invasive vegetation have a 89.62% chance of being smaller than those recorded at sites with native vegetation (mean difference of -0.71 ±0.88 m/hour, Cr.I. 95% -2.62–0.87 m/hour; Fig. 5B).

**Fig. 5.**
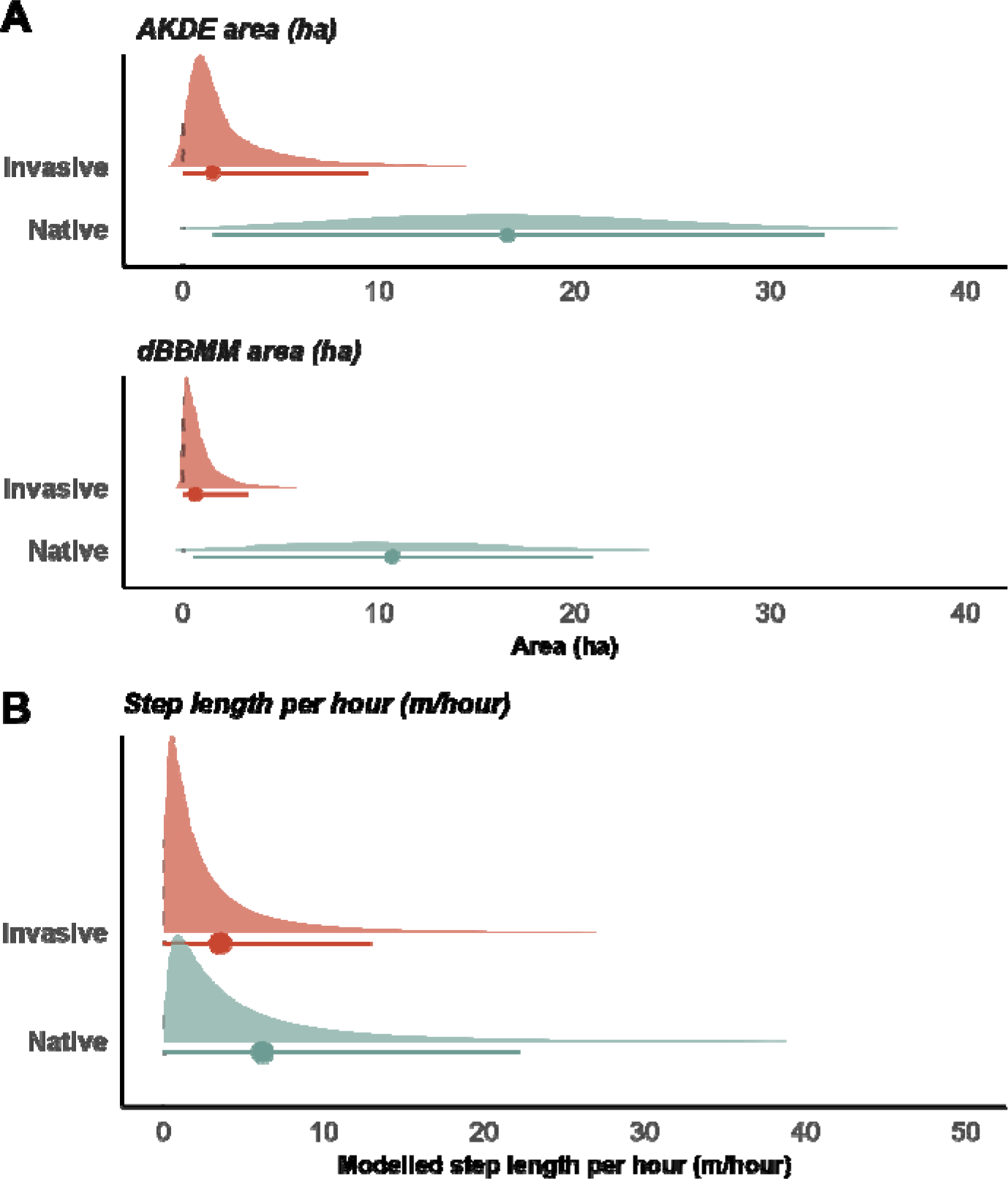
Posterior distributions associated with vegetation effects on western tiger snake (*Notechis scutatus occidentalis*) space use and movements. Sites with invasive vegetation: Herdsman Lake and Kogolup Lake, those with predominantly native vegetation: and Yanchep National Park and Black Swan Lake. A) Posterior distribution resulting from Bayesian comparisons of space use. Space use measures used were point estimates of 95% utilisation distribution contours from autocorrelated kernel density estimators (AKDE) and the 95% occurrence distribution contours from dynamic Brownian bridge movement models (dBBMM; both in hectares). B) Posterior distribution of population effect of vegetation on within day step lengths per hour. Dots indicate point estimates, and lines indicate the 95% median HDI credible intervals.

## DISCUSSION

We observed clear differences in estimates of overall space use for male tiger snakes occupying wetlands dominated by invasive kikuyu grass compared to those occupying wetlands with native vegetation, with snakes using smaller areas at sites with kikuyu. However, there was considerable variation in within-day step lengths per hour between individual snakes, resulting in substantial overlap for within-day step lengths per hour across sites.

Snakes in wetlands dominated by kikuyu grass spent most of their time within the invasive grass, covering overall smaller areas within the two month study period. A suite of factors influence intra- specific variation in home range and movement, such as habitat composition and fragmentation, resource availability, weather conditions, sex, and age (Rivrud et al. 2010; Braham et al. 2015; Mayer et al. 2019). It is not uncommon for animals in urban and other environments with high human impacts to have smaller home ranges than those living in rural or less-disturbed environments (Lowry et al. 2013; Tucker et al. 2018). For mammalian species persisting in areas with both high and low human impact these reduced movements can result from both movement barriers and from increased resource availability (Tucker et al. 2018) with smaller home ranges and shorter movements associated with better habitat quality (Fustec et al. 2001; Bjørneraas et al. 2012). The king cobra (*Ophiophagus hannah*), another elapid snake species, had reduced movement in human altered landscapes being restrained to small areas of relatively natural remnant vegetation in agricultural environments compared with protected forest areas (Marshall et al. 2020). For cobras, movement barriers appear responsible for reduced movements in agricultural landscapes. Similarly, Maddalena et al. (2020) reported that home ranges of milksnakes (*Lampropeltis triangulum*) were smaller in urban parks compared to a more natural study site, likely due to fragmentation by roads and differences in habitat composition.

For tiger snakes, sites dominated by kikuyu and native vegetation did not differ substantially with respect to prey availability, temperature or predation pressure for tiger snakes (Cornelis et al. 2022), so we suspect these factors are not directly driving the observed differences in tiger snake movements.

The major habitat difference between the vegetation types is that kikuyu forms more structurally dense cover, completely covering the ground (compared to patches of bare ground which occur within native vegetation). We suspect this structural difference allows tiger snakes to move vertically between basking locations on top of kikuyu and shelter sites at the base, rather than having to travel horizontally along the ground to open basking areas in sites dominated by native vegetation (JC & DCL pers obs). Consequently, tiger snakes living in kikuyu-dominated habitats have smaller home ranges than those in native vegetation reflecting their predominantly vertical rather than horizontal movements due to increased density of resources in the form of basking and shelter sites. For some reptiles, invasive plants which provide structurally complex habitats can be important for their persistence in highly modified environments. Garden et al. (2007) reported a positive association between native reptiles and up to 50% weed cover in urban remnant habitat fragments. For these species it is the habitat structure, rather than the plant species composition, that drives reptile persistence.

Another explanation for the smaller areas used by snakes in kikuyu-dominated habitats could be a smaller potential area for them to occupy. The two wetlands with kikuyu are surrounded by historic urbanisation and roads, especially HL, which has been subject to anthropogenic modification since the 1850’s (Clarke et al. 1990; Gentilli and Bekle. 1993; Kelobonye et al. 2019; Lettoof et al. 2021b). Some connectivity remains between KL and adjacent wetlands as part of the broader Beeliar Regional Park yet multiple roads are interspersed through this region which act as a habitat barrier and can lead to increased mortality (Andrews and Gibbons 2005; Lettoof et al. 2021b; Cornelis et al. 2021). Roads are well known for inhibiting movements of urban wildlife (Clark et al. 2010, Holderegger & Di Giulio 2010, Doherty et al. 2020) and this, along with habitat fragmentation, may exert selection pressure against extensive movement of snakes at HL and KL. At BS, where surrounding urbanisation is relatively recent (< 30 years) and the main road separating the lake from adjacent wetlands was only developed in 2010 (Google Earth 2021), one snake left the wetland and spent time around industrial infrastructure and another spent much of its time moving along the edge of a main road.

These behaviours appear to be an attempt to reach a neighbouring southern wetland; the shorter period of isolation may have been insufficient to induce a selective pressure for this population to avoid risky urban environments (Shepard et al. 2008).

Our findings add to an increased understanding of how changes in habitat composition, driven by invasive vegetation, can affect space use by animals. Our current findings suggest that management plans for urban wetlands in the Perth urban area should improve structural complexity of homogenous kikuyu habitat by planting native species to support the predators in these ecosystems (Cornelis et al. 2022). However, it remains to be determined if and how this grass affects reproductive success and long-term survival of tiger snakes. For example, it is possible that apart from the apparently positive effect of the increased structural complexity of invasive grass for thermoregulatory behaviour, changes in vegetation structure might negatively impact the foraging efficiency of tiger snakes. In other systems, increased structural complexity of the habitat decreases predation efficiency (e.g. Warfe & Barmuta 2004), although for snakes increased vegetation complexity can improve predation efficiency (e.g. Koenig et al. 2007; Somsiri et al. 2020). Moreover, our study was restricted to a comparatively short period and to mature males. Longer-term studies including both sexes could investigate the role of habitat homogenisation via invasive vegetation across seasons and depending on sex, as females might have different habitat requirements than males, especially during the reproductive season (Brown et al. 2002).

## DATA ACCESSIBILITY

Data used in this study is avaible on Zenodo (DOI: https://doi.org/10.5281/zenodo.7700983). The Zenodo repository also includes all R scripts used to run analyses and generate figures.

## ACKNOWLEDGEMENTS

We thank Katrina Wood for volunteering her time to provide advice and practical training concerning snake surgeries, Aleesha Turner for volunteering her time to assist with surgeries, husbandry and tracking of tiger snakes for this project, and Bill Bateman for allowing us to use his lab for snake husbandry. We also pay respects to the traditional owners of the land, the Whadjuk-Noongar people, where this research was conducted. The use of wild animals for this project followed the Australian Code of Practice for the care and use of animals for scientific purposes and was approved by the Curtin University Animal Ethics committee (ARE2020-06). We also thank the staff at the Department of Biodiversity Conservation and Attractions and the staff at Regional Parks for allowing us access to the study sites and issuing the required permit (FO25000294-2).

## AUTHOR CONTRIBUTIONS

All authors contributed to the study conception and design. Surgical procedures were performed by Christine Cooper and Jari Cornelis. Material preparation and data collection were performed by Jari Cornelis and Damian Lettoof. Data analysis was performed by Benjamin Marshall. The first draft of the manuscript was written by Jari Cornelis and Benjamin Marshall and all authors commented and contributed to subsequent versions of the manuscript. VHF transmitters were provided by Martin Mayer.

