## Supplementary material for "Stuck in the weeds: Invasive grasses reduce tiger snake movement"

Table S1. Summary of Autocorrelated Kernel Density Estimators (AKDE) results from the top fitting models. Including point estimates from 95% contours (ha) and associated 95% confidence intervals, the effective sample size (DOF_area), the type of top fitting model (Model abbreviations: Ornstein-Uhlenbeck [OU], Ornstein–Uhlenbeck Foraging [OUF]), the home range crossing time (days; tpositionEst), Dynamic Brownian Bridge Movement Models (dBBMM) 95% confidence areas (ha).

| **Snake ID** | **aKDE_low** | **aKDE_est** | **aKDE_high** | **DOF_area** | **Movement_model** | **tpositionEst** | **dBBMM_area** |
|---|---|---|---|---|---|---|---|
| HL162 | 0.40 | 0.60 | 0.83 | 29.26 | OUF anisotropic | 2.75 | 3.11 |
| HL166 | 0.08 | 0.12 | 0.15 | 37.80 | OU anisotropic | 3.05 | 0.03 |
| HL168 | 0.17 | 1.11 | 2.92 | 2.32 | OU isotropic | 16.51 | 0.08 |
| HL88 | 18.85 | 234.05 | 713.27 | 1.59 | OUF anisotropic | 1.97 | 0.62 |
| KL01 | 0.93 | 1.60 | 2.44 | 16.94 | OUF anisotropic | 5.62 | 0.09 |
| KL02 | 0.38 | 0.61 | 0.89 | 22.38 | OUF anisotropic | 4.49 | 0.05 |
| KL06 | 0.72 | 12.90 | 42.10 | 1.34 | OU anisotropic | 1.89 | 0.03 |
| BS01 | 14.23 | 34.24 | 62.90 | 7.43 | OU anisotropic | 4.81 | 30.62 |
| BS02 | 1.33 | 6.56 | 15.86 | 2.96 | OU anisotropic | 15.97 | 0.61 |
| BS08 | 9.62 | 21.48 | 38.03 | 8.61 | OU anisotropic | 3.00 | 17.15 |
| BS09 | 19.11 | 130.30 | 345.96 | 2.28 | OU anisotropic | 17.84 | 25.65 |
| Y58 | 0.51 | 1.28 | 2.39 | 6.91 | OU isotropic | 2.31 | 0.21 |
| Y63 | 7.45 | 24.44 | 51.35 | 4.59 | OUF anisotropic | 7.15 | 2.59 |
| Y65 | 3.93 | 8.95 | 16.00 | 8.27 | OU anisotropic | 4.57 | 4.97 |

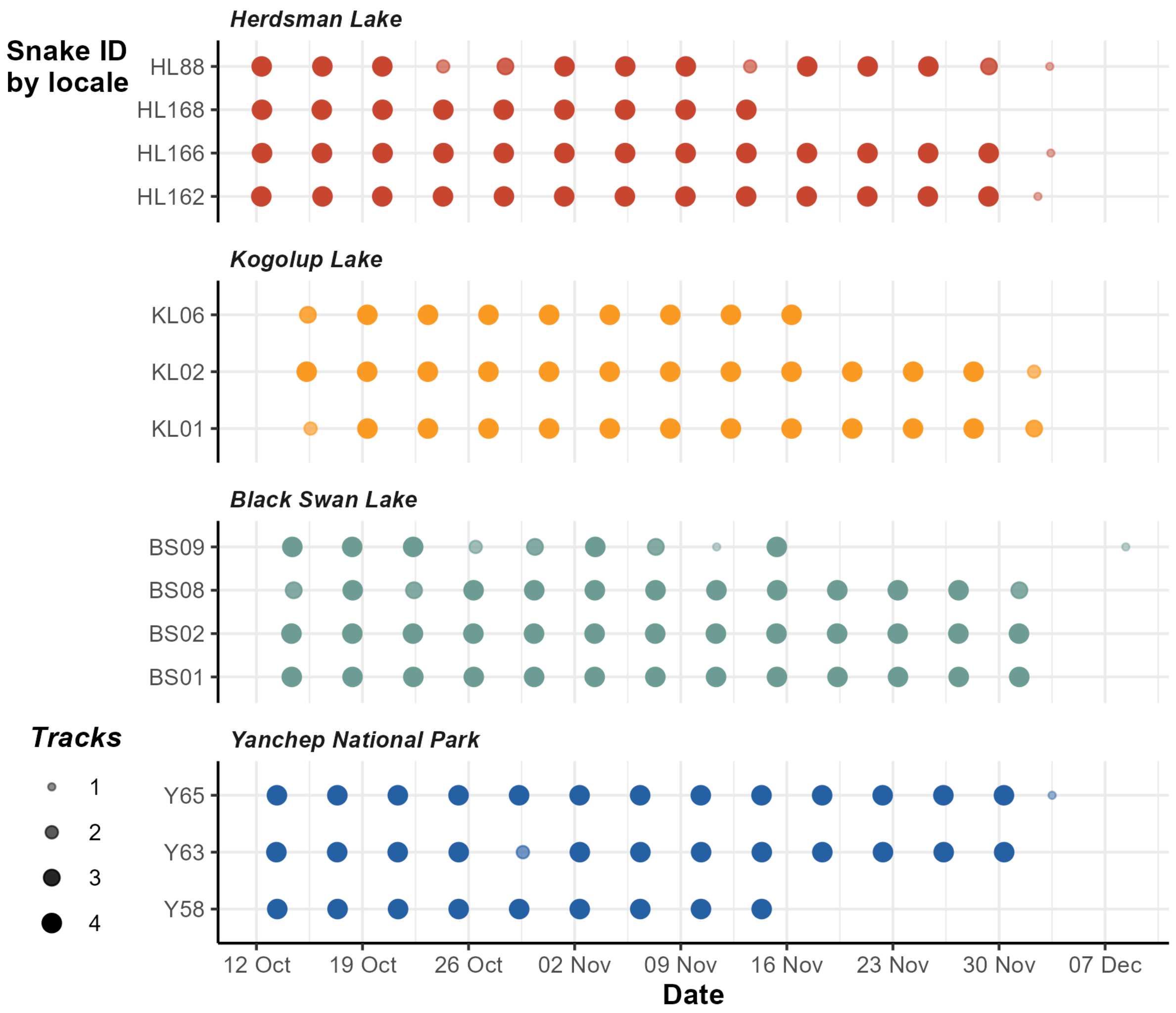


Fig. S1. Radio-tracking dates and number of locations successfully recorded per day for individual tiger snakes (*Notechis scutatus occidentalis*) studied at four wetlands (Herdsman Lake, Kogolup Lake, Black Swan Lake and Yanchep National Park) in the Perth metropolitan area, Western Australia.

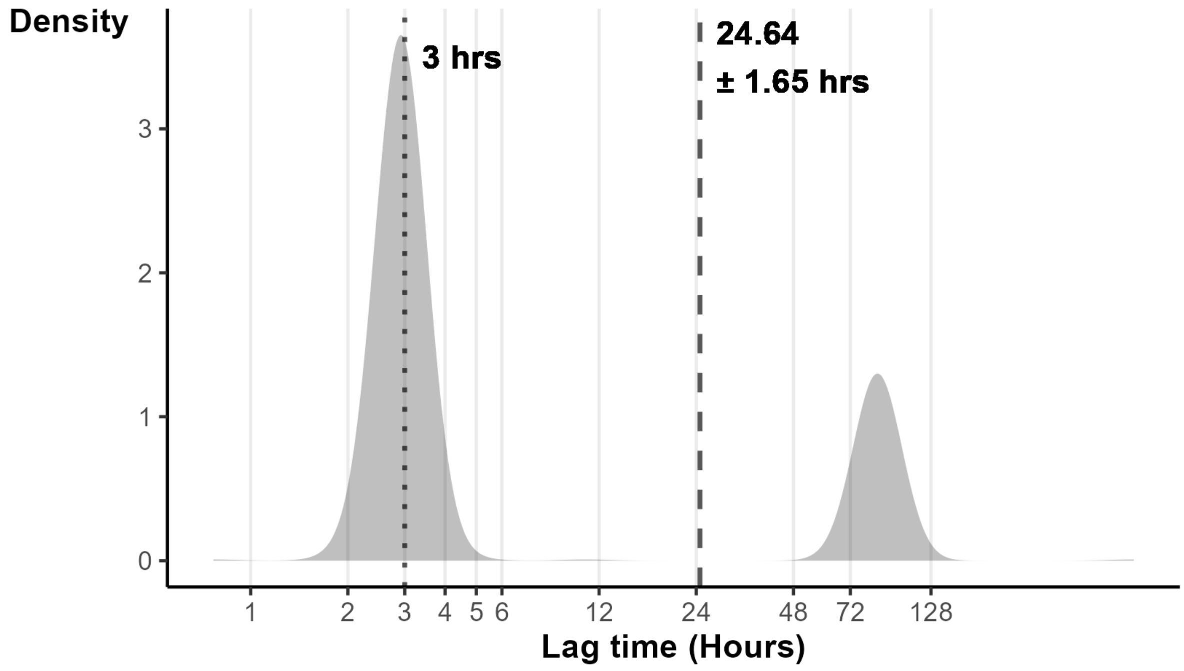


Fig. S2. Density plot showing the time lag in hours between tracks, with median (dotted) and mean ± standard error (dashed) annotated, for tiger snakes (*Notechis scutatus occidentalis*) radio-tracked at wetlands in the Perth metropolitan area, Western Australia. Snakes were radio-tracked every four days, and were located up to four times per day at intervals of approximately 3 hours.

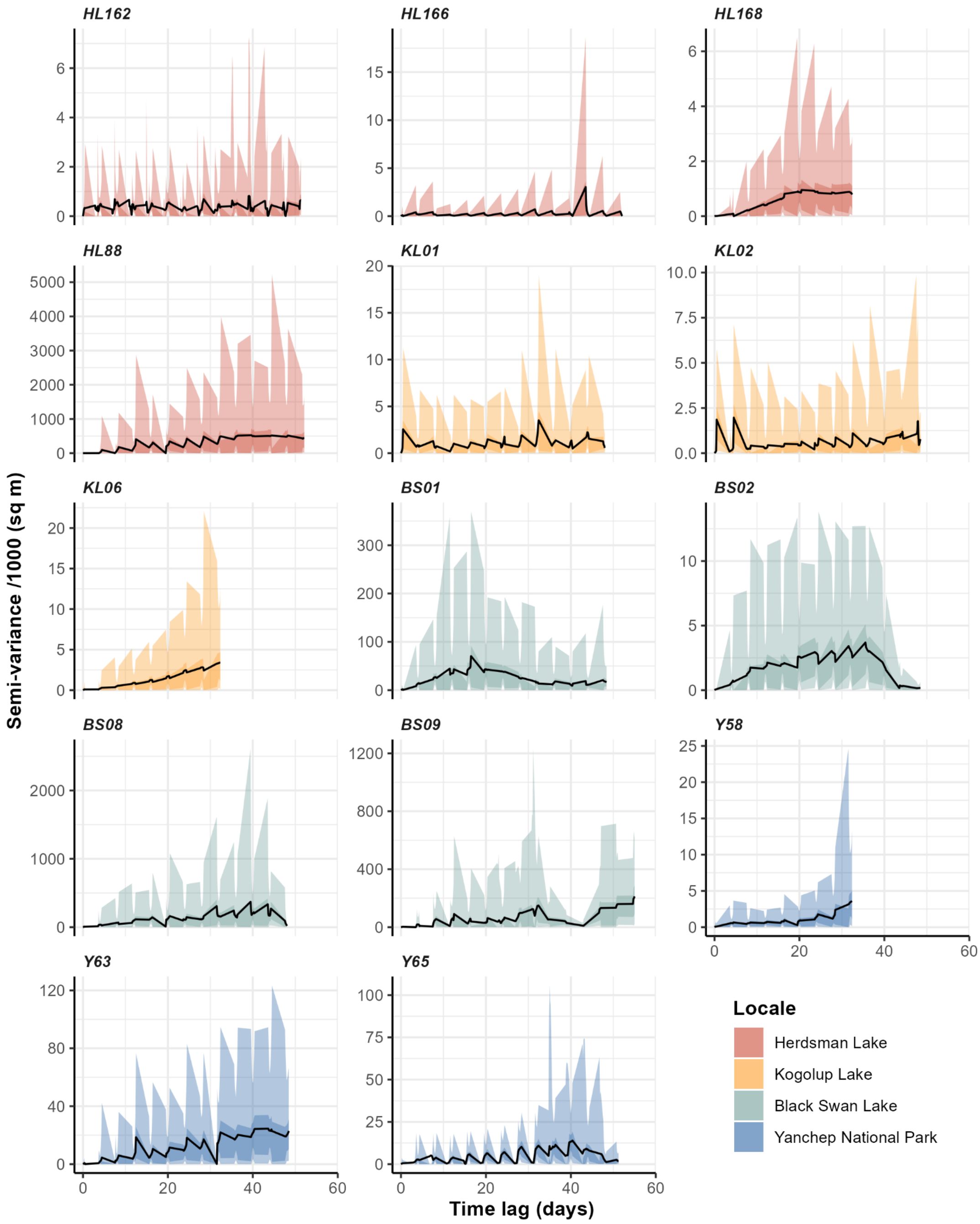


Fig. S3. Variogram results displaying uncertainty in the semi-variance of home range area estimates with x-axis starting at the beginning of each individual's tracking period. Shaded areas display the 50% (dark shading) and 95% (light shading) confidence intervals for all individual tiger tiger snakes (*Notechis scutatus occidentalis*) radio-tracked at four wetlands in the Perth metropolitan area, Western Australia. Warmer colours (Herdsman Lake and Kogolup Lake) are the locales with invasive grass; cooler colours (Black Swan Lake and Yanchep National Park) are the locales with native vegetation.

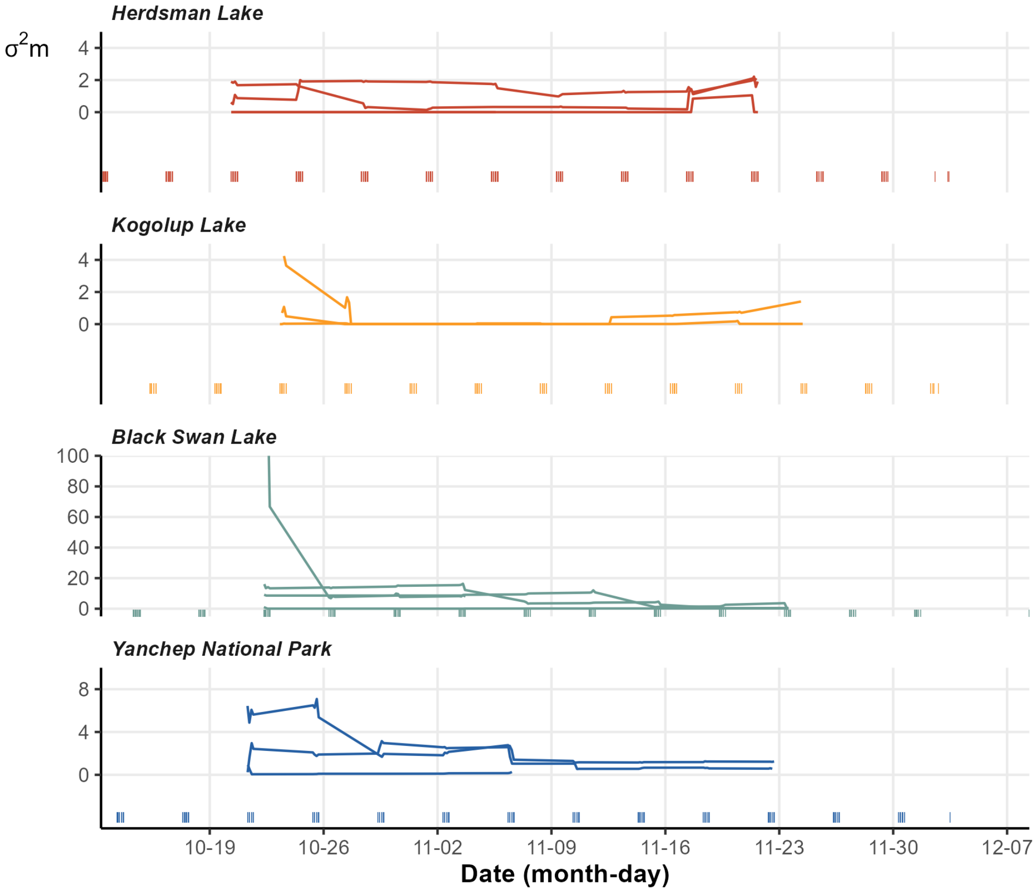


Fig. S4. Motion variance extracted from Dynamic Brownian Bridge Movement models showing the movement capacity calculated for tiger snakes (*Notechis scutatus occidentalis*) radio-tracked at four wetlands in the Perth metropolitan area, Western Australia. Warmer colours (Herdsman Lake and Kogolup Lake) are the locales with invasive grass; cooler colours (Black Swan Lake and Yanchep National Park) are the locales with native vegetation. $\sigma^2$m represents a proxy for movement capacity.
